# Network analysis reveals converging molecular mechanisms in genetic epilepsies

**DOI:** 10.64898/2026.10.28.762494

**Authors:** Yuanyuan Duan, Alexandra Valeanu, Rob P.W. Rouhl, R. Jeroen Vermeulen, Theo M. de Kok, Friederike Ehrhart

**Author notes:** Corresponding author; Yuanyuan Duan. Alexandra Valeanu; Rob Rouhl; Jeroen Vermeulen; Theo M. de Kok; Friederike Ehrhart.

## Abstract

**Background:** Rare genetic epilepsies are often difficult to diagnose and treat. They can arise from changes in many different genes, and the molecular basis linking these changes to epilepsy remains poorly understood. This complexity makes it difficult to select effective treatments and develop new therapies, leaving many patients with poorly controlled seizures and reduced quality of life

**Methods:** To address this gap, we first identified epilepsy-associated genes and conducted a network-based analysis of biomolecular interactions in genetic epilepsy to identify converging pathways with high translational relevance and potential therapeutic targets. To run the analysis, we constructed a human brain-specific molecular interaction network by integrating all human pathways from the WikiPathways database. HotNet2 was then applied to identify highly perturbed clusters through evidence-based weight diffusion of EAGs in the human brain network.

**Results:** We identified 2,496 epilepsy-associated genes (EAGs) from multiple resources. These genes encode coding for metabolite interconversion enzymes (20%), transporters (15%), and protein-modifying enzymes (11%). Furthermore, the network analysis identified 19 subnetworks with more than two nodes and 125 unique genes. Gene enrichment analysis revealed convergence of processes including action potential, synaptic function, nervous system development, TOR signaling, cilium organization, DNA repair, and cell fate specification. We identified 44 first-order neighbors connecting the subnetworks. By integrating drug-target interactions from DrugBank using CyTargetLinker, we highlighted potential therapeutic targets within the S4 subnetwork, associated with cerebral cortex development, as well as CREB1, a connector between subnetworks.

**Conclusion:** Network analysis organized heterogeneous epilepsy-associated genes into biologically related subnetworks and generated hypotheses for further research.

## INTRODUCTION

Epilepsy is a common, chronic neurological condition characterized by recurrent, unprovoked seizures (1). It has diverse etiologies, including developmental brain abnormalities, infection, traumatic brain injury, stroke, brain tumors, and genetic factors; however, some underlying causes remain unclear (2).

Genetic variation is thought to explain up to two-thirds of the overall liability for epilepsy in the general population, and genetic susceptibility can be attributed to monogenic, polygenic, and other multifactorial factors (3,4). Although epilepsy is not rare, many monogenic forms are rare. For instance, rare genetic epilepsies include Angelman syndrome, linked to *UBE3A* (5) and Dravet syndrome, mostly caused by *SCN1A* mutations (6). Genetic epilepsy exhibits a wide range of clinical phenotypes associated with heterogeneous genotypes (7,8).

Integrating biomolecular interactions relevant to disease manifestations can improve understanding of the relationships among biological entities and the mechanisms of information flow in complex biological systems. This approach has helped elucidate disease mechanisms, as demonstrated in studies on Rett syndrome (11) and 22q11.2 deletion syndrome (12). One application of systems biology is to identify disease-relevant subnetworks within these large and complex molecular networks (9,10). These subnetworks can help guide disease mechanism summary and drug targeting development.

Various analytical approaches can be used to explain biology from large-scale networks, each with its own added value. Among these, heat diffusion provides a unique perspective to identify network regions in a large network. It models the spread of a biological perturbation by assigning an initial "heat" to seed nodes and allowing the signal to propagate through the protein–protein interaction (PPI) network. This framework is biologically motivated by the observation that disease phenotypes typically arise from dysregulation of interconnected clusters of genes rather than isolated genes (13,14).

HotNet2, a heat diffusion-based algorithm, employs a directed heat diffusion model that evaluates both the attributes of individual genes (with high biological importance) and the local network architecture of their neighboring interactions, thereby addressing the shortcomings inherent in traditional pathway enrichment analyses and topology-based network methods (15). It has been shown to be effective in identifying disease-associated genes, including its use in the *Mycobacterium tuberculosis* PPI network with genome-wide association study (GWAS)-derived mutation significance scores (16) and in detecting gene subnetworks associated with complex traits and diseases (17). A diffusion-based network analysis approach may help identify disease pathways and previously overlooked molecular effectors that contribute to seizure manifestations through direct or indirect modulation of neuronal excitability and inhibition.

This study aims to construct a comprehensive biomolecular interaction network by integrating human pathways from WikiPathways (WP) to represent the molecular mechanisms underlying human biology (18). We assigned heat scores to epilepsy-associated genes and propagated the heat through the network. Clusters of interacting proteins that collectively retain high heat levels could be associated with the pathogenesis of genetic epilepsy. We further investigated these subnetworks to identify potential modulators of disease mechanisms and to evaluate their potential as targets for drug-based therapy. This approach could offer novel therapeutic targets that modulate the balance between neuronal excitation and inhibition and expand the landscape of treatment opportunities for genetic epilepsy.

## METHODS

### Epilepsy-associated gene curation

Genes were retrieved from the following publicly available human genotype-phenotype databases: Online Mendelian Inheritance in Man (OMIM) (19), DisGeNET (20), ClinVar (21), and Orphadata (22). Searches used the following keywords: *epilepsy, epileptic, epilepsies, seizures, seizure, convulsion, and convulsions*. The data were collected in June 2026. Database-specific selection criteria were used to identify genes strongly associated with epilepsy. In OMIM, the genes were retained only if the molecular basis was known or if contiguous gene duplication or deletion syndrome was involved (phenotype mapping key ≥ 3). In ClinVar, only the variants classified as pathogenic or likely pathogenic were retained. In DisGeNET, genes with only curated or literature-derived disease associations (gene-disease association score ≥ 0.4) were included with academic access. From Orphanet, we identified the Human Phenotype Ontology (HPO) (23) terms with phenotype keywords and retrieved the associated genes from Orphadata (release dated 23 Jun 2026). Gene–phenotype associations with reported phenotype frequencies above 30% were included. Gene identifiers obtained from the four resources were mapped to Ensembl gene IDs using the MyGene Python package (version 3.2.2) (24). We used the resulting curated genes, termed epilepsy associated genes (EAGs), as the primary dataset for downstream analysis.

### Functional enrichment and pathway overrepresentation analysis

Functional enrichment and pathway overrepresentation analyses were performed using g:Profiler (25). We submitted Ensembl IDs as input and restricted the organism to *Homo sapiens*. All human genes were used for background. The analysis included the molecular function (MF), cellular component (CC), and biological process (BP) categories of Gene Ontology (GO; release dated 2026-01-23) (26) and WikiPathways (WP; version 20260310) (18). P-values were adjusted for multiple testing using the false discovery rate (FDR) method, with an adjusted p-value threshold of 0.05. We also annotated the EAGs using the PANTHER Protein Class classification system (27) to categorize their encoded proteins by function.

### Biomolecular interaction network construction

Biomolecular interactions were collected from WP (version 20260810) and used as a signaling network for diffusion analysis. All direct human interactions were retrieved using SPARQL endpoints. Gene product identifiers were mapped to Ensembl IDs to harmonize identifiers using the MyGene API (24). We then filtered genes expressed at nTPM (normalized transcript expression per million) ≥ 1 in at least one brain region in the Human Protein Atlas (HPA) (28). To reduce the overrepresentation of ubiquitous biological molecules, 191 common small molecules such as H⁺, H₂O, and ATP were removed (11). Duplicate edges and self-loops were excluded from analysis. All interactions were represented as source and target and integrated into one network.

### HotNet2 Clustering

We applied the HotNet2 algorithm (15), which combines attribute-based and topology-driven clustering to identify significantly perturbed subnetworks by input heat that are unlikely to occur by chance. As input for HotNet2, two files were required: (i) a heat-score vector assigning a numerical value to each node, and (ii) an edge list describing network connectivity. Heat scores were defined based on evidence from four sources (OMIM, Orphadata, ClinVar, and DisGeNET) expressed as counts from 1-4. Nodes that were not classified as EAGs received no heat.

The algorithm depends on two parameters: insulation parameter *β* and subnetwork threshold *δ*. We set *β* = 0.5 by default, allowing half of the heat to diffuse to neighboring nodes at each iteration. The *δ* threshold was empirically determined using 100 permutation statistical tests.

We followed the implementation workflow available at: https://jhrcook.github.io/HotNetvieweR/articles/HotNet2-setup-and-example.html#setup.

### Drug identification

Approved drugs for epilepsy (MONDO:0005027) were retrieved from the Open Targets Platform (29). We used CyTargetLinker to identify drug-targeting subnetwork components from DrugBank (version 4.2) (30).

## RESULTS

### Epilepsy-associated genes

From four human phenotype-genotype databases, including OMIM, Orphadata, DisGeNET, and ClinVar, we obtained records associated with epilepsy, seizures, and convulsions (**Supplementary File 1, Tables S1-S4**). All retrieved records were mapped to Ensembl IDs for identification (**Supplementary File 2, Fig S1**). We retained only the Ensembl-identified genes for further analysis. The complete list of genes is provided in **Supplementary File 1, Table S5**.

In total, 2,496 unique genes were associated with epileptic phenotypes (**Fig 1A**). Of these, OMIM contributed 2,136 genes, representing the largest portion of the dataset. Of these OMIM genes, 1,349 (63%) were unique to OMIM, whereas 787 (37%) overlapped with the other three databases. Orphanet contributed 811 genes, DisGeNET 508 genes, and ClinVar 74 genes. Only 35 genes (1.4%) overlapped across the four databases.

**Fig 1.**
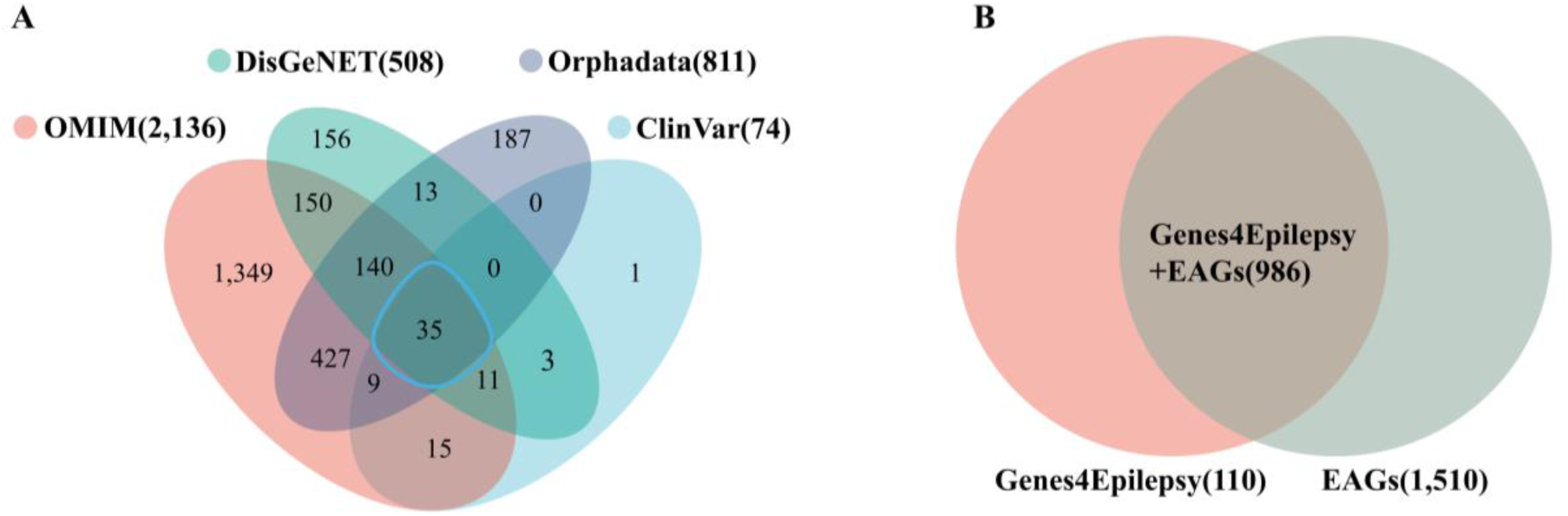
EAG coverage across four phenotype-genotype databases and cross-referenced with the Genes4Epilepsy database. **(A)** Venn diagram of the distribution of EAGs from ClinVar, Orphadata, DisGeNET, and OMIM. **(B)** Venn diagram of the overlap of EAGs and Genes4Epilepsy.

To validate our curated gene list, we cross-referenced the dataset with the Genes4Epilepsy database (n = 1,096) (v2026-09) (31) (**Fig 1B**). We identified 986 overlapping genes, representing 90% (986/1,096) of the Genes4Epilepsy genes. These results indicate that the EAG curation captured the majority of genes included in Genes4Epilepsy while expanding the collection with additional phenotype-associated genes.

### Functional characterization and pathway overrepresentation

Of all the EAGs, 1,996 (80%) genes were mapped to their corresponding proteins and classified into protein classes in PANTHER (**Fig 2**). The most prevalent categories were metabolite interconversion enzymes (20%), transporters (15%), protein-modifying enzymes (11%), and gene-specific transcriptional regulators (8%). The significantly enriched terms from GO and WP (adjusted p-value < 0.05) are listed in **Supplementary File 1, Table S6**. The top ten enriched terms in GO MF and GO CC and overrepresented pathways in WP were visualized in **Supplementary File 2, Fig S2A-C**. The top ten most significantly enriched GO BP terms ranked by the adjusted p-values are shown in **Fig 2B**. In GO BP, *system development* (p_adj_ = 2.7e^-102^) was the most significantly enriched, followed by *nervous system development* (p_adj_ = 4.1e^-100^) and *multicellular organism development* (p_adj_ = 7.8e^-100^). These enriched terms highlight the broad involvement of developmental processes in genetic epilepsy pathogenesis.

**Fig 2.**
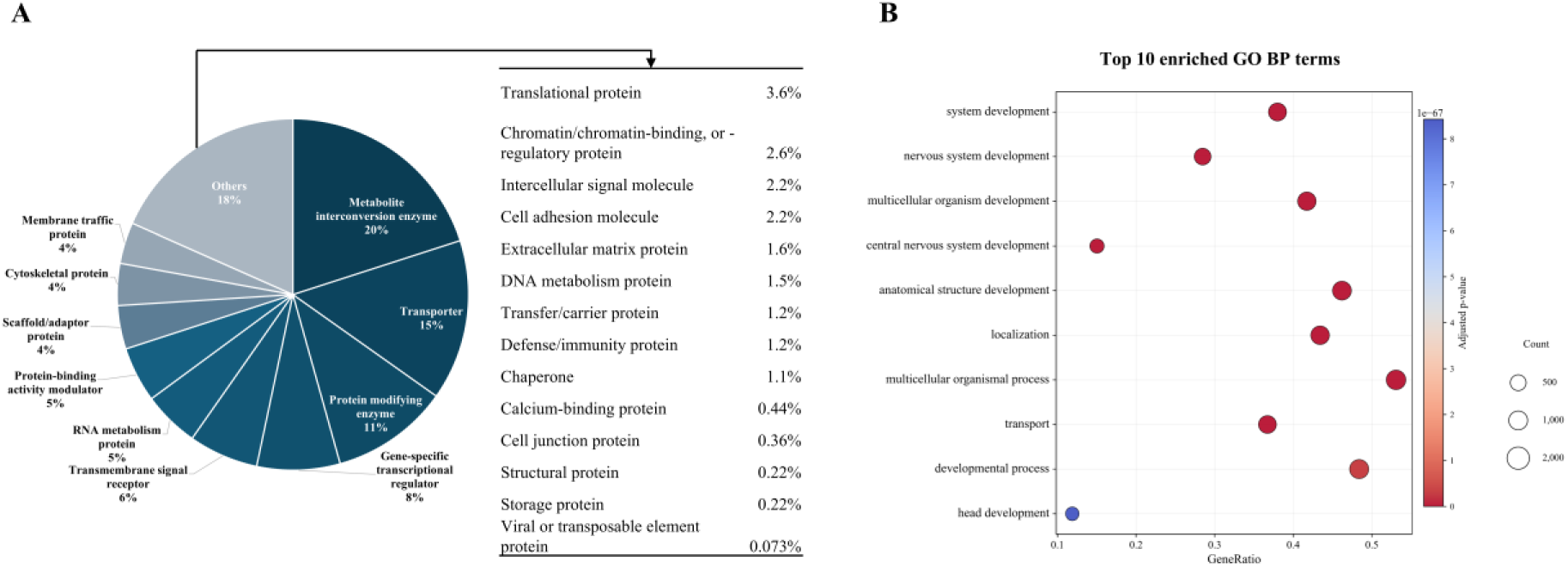
**(A)** Protein class distribution of EAGs using PANTHER. (B) Gene enrichment analysis sourced from GO BP. The top ten enriched terms are illustrated and ranked by padj.

### Biomolecular interaction network from WP

To build a brain-specific molecular network, we integrated interactions from 1,016 human pathways from the WP. A total of 19,421 unique undirected edges were retrieved (**Supplementary File 2, Fig S3).** Genes that could not be mapped to Ensembl IDs or were not expressed in the brain, and common metabolites were excluded. The final network comprised 11,955 unique undirected edges, 3,117 unique gene products, and 4,331 metabolite nodes (**Supplementary File 1, Table S7**). After integrating the interactions into a single molecular interaction network, the nodes were ranked by degree. The top ten degree-ranked nodes in descending order are AKT1, TP53, MAPK14, CREB1, STAT3, CTNNB1, NFKB1, MAPK1, SRC, and KDR. Reactive oxygen species (ROS, CHEBI:26523) ranked 11^th^ and first among all metabolites. The most frequent interaction types were annotated as direct interaction (n = 4,550), conversion (n = 3,172), stimulation (n = 1,172), and inhibition (n = 1,040).

### Heat selection and parameter assessment

We assigned a weight to each node based on its supporting evidence strength as an EAG across the four databases. To assess the effect of the weight design, we first compared four weighting schemes: cardinal weights (0, 1, 2, 3, and 4), scaled weights (0, 10, 20, 30, and 40), binary weights (0 or 1), and randomly assigned weights to the EAG nodes (0, 1, 2, 3, and 4). The integrated network defines the paths for heat diffusion, with node weights serving as the initial heat. We compared the Jaccard similarities across the four weight schemes after propagation to determine the weight choice (**Supplementary File 2, Fig S4**). The cardinal and scaled weighting schemes produced identical results, indicating that the proportional rescaling of the heat scores did not alter the subnetwork output. Binary weights yielded a Jaccard similarity of

0.54 relative to cardinal weights, whereas random weights produced outputs that differed more from those obtained using both evidence-based schemes (binary and cardinal). These comparisons indicate that the subnetwork identification is sensitive to the initial heat. We retained the source count weights to preserve perturbation differences in the database support. HotNet2 generated four candidate δ thresholds representing different levels of edge stringency (**Supplementary File 1, Table S8**). Higher δ values retain stronger connections in the heat-diffusion graph, resulting in smaller subnetworks. At the lowest threshold (δ = 0.0176), the largest subnetwork contained 156 nodes, whereas at the highest threshold (δ = 0.04547), large subnetworks were fragmented into smaller subnetworks (**Supplementary File 2, Fig S5**). Therefore, for biological interpretation, we chose the highest δ threshold for subsequent module-level analyses, as it resembles manageable subnetwork sizes for visualization and biological interpretation. At this threshold, HotNet2 identified 51 subnetworks containing at least two genes, comprising 189 unique genes. Nineteen subnetworks contained at least three genes (125 genes), and ten contained at least five genes (95 genes).

To assess the statistical significance of the output, we generated 100 test networks with randomized edge swaps while preserving their degrees. At the highest δ, the observed number of subnetwork sizes (≥ 2 nodes) was more than expected by permutation (p-value ≤ 0.01) (**Supplementary File 2, Fig S6**).

### Subnetwork visualization and their biological annotation

The 10 subnetworks (n >= 5) were visualized and annotated with the most significant BP terms in **Fig 3**. S1 (*nervous system development*) was the largest module, followed by S2 (*cardiac muscle cell action potential*), S3 (*chromatin organization*), and S4 (*cerebral cortex development*). Two modules were associated with synaptic functions: S6 (*chemical synaptic transmission*) and S10 (*cynaptic vesicle exocytosis*). The other subnetworks are S5 (*TOR signaling*), S7 (*cell fate specification*), S8 (*cilium organization*), and S9 (*DNA repair*). We observed that all the identified components were EAGs.

**Fig 3.**
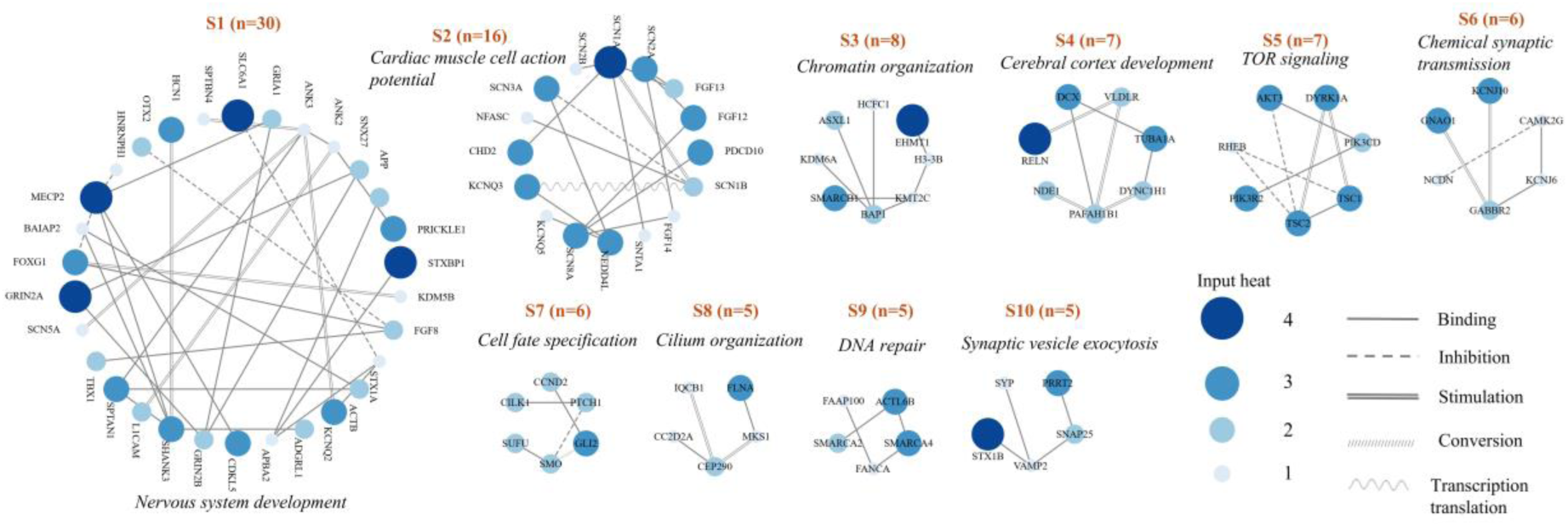
Ten distinct subnetworks (n >= 5) identified from the WP signaling pathway at the highest δ using HotNet2. The node size and color density indicate the initial evidence score related to being associated with epilepsy.

We also explored whether some subnetworks were interconnected by their first neighbors. By comparing the overlapping first neighbors of each subnetwork, we found that 44 nodes connected the subnetworks (**Supplementary File 1, Table S9**). According to the Human Transcription Factors Database (32), 15 of these are transcription factors. Among them, CREB1 (cAMP response element-binding protein 1) connects the most subnetworks (S1, S5, S11, S12 and S41). Interestingly, of the ten highest-degree nodes from the original network, six were among the 44 connectors. Collectively, the results indicate that the high-degree nodes tended to be connectors, and high-heat nodes tended to be subnetwork components.

### Clinical relevance and drug repurposing exploration

To further assess the relevance of the identified subnetworks to epilepsy, we compared the modular components with approved ASM (antiseizure medication) targets from Open Targets (**Supplementary File 1, Table S10**). Of the 93 ASM targets, 20 (21%) were present in the identified modules (**Fig 4A**). The targets included sodium channel subunits (SCN1A, SCN2A, SCN3A, SCN5A, and SCN8A), gamma-aminobutyric acid receptors (GABRA1, GABRA2, GABRA3, GABRB3, and GABRD) and the transporter SLC6A1, glutamate receptor subunits (GRIN1, GRIN2A, GRIN2B, GRIA1, GRIA3, and GRIA4), and potassium channel subunits (KCNQ2/3/5). We conducted a hypergeometric test to determine whether the ASM targets were significantly enriched compared to the input of 3,117 gene products in the background. In total, 32 targets were present in the background network. We found a 10.31-fold enrichment relative to the background (*P* = 2.01 × 10⁻¹^7^). This overlap supports the high relevance of subnetworks in establishing therapeutic targets in epilepsy.

**Fig 4.**
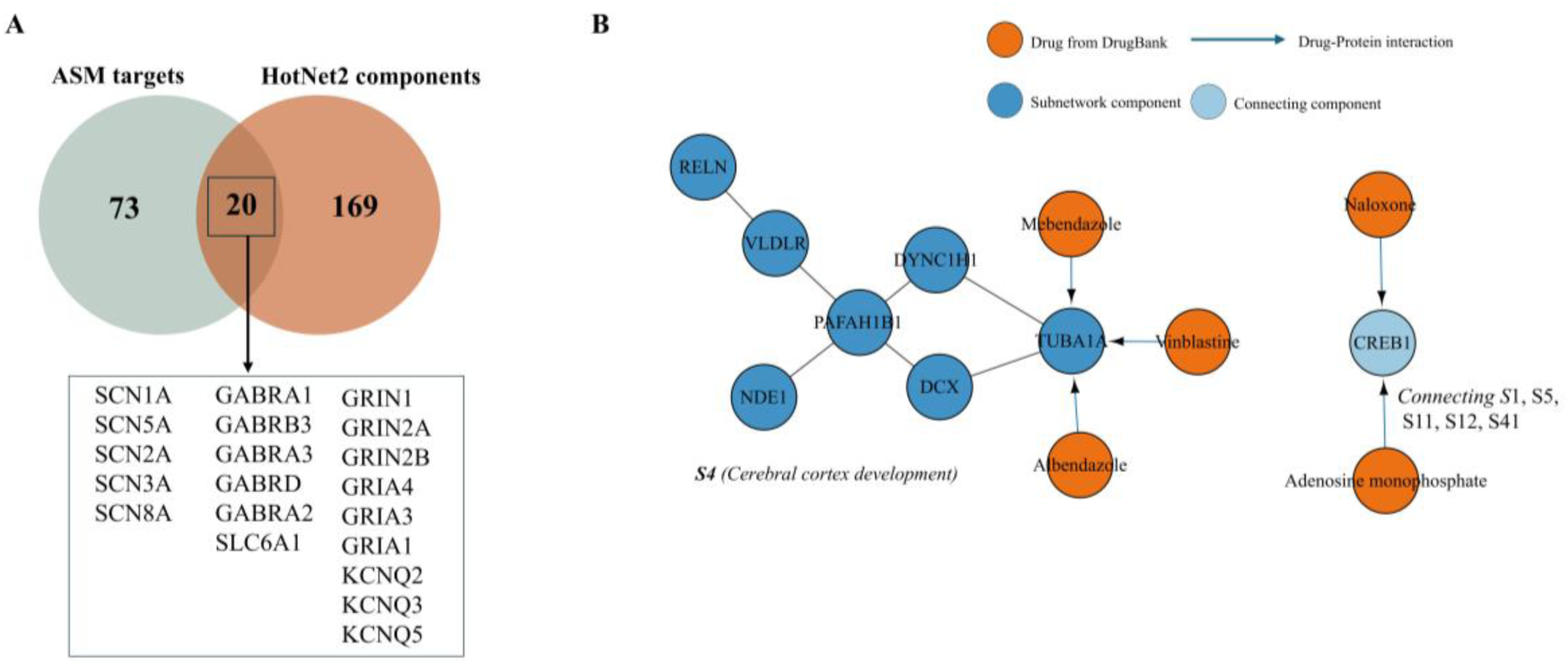
**(A)** Comparison of ASM targets (n = 93) from Open Targets with all HotNet2-identified components (n = 189 when including all unique genes identified). **(B)** S4 and CREB1 extended with approved drugs from DrugBank.

The remaining components from the individual subnetworks can be assessed for possible drug repurposing candidacy in genetic epilepsy. We used CyTargetLinker to extend the subnetworks with curated drug-target interaction knowledge from DrugBank. We extended S4 (*cerebral cortex development*) and CREB1, the subnetwork connector (**Fig 4B**). From S4, we identified three drugs (mebendazole, vinblastine, and albendazole) targeting TUBA1A. CREB1 is targeted by naloxone and adenosine monophosphate.

## DISCUSSION

In recent decades, advances in sequencing technologies have substantially expanded our understanding of the genetic basis of epilepsy (33). Many epilepsy-associated genes encoding proteins are not directly involved in neuronal excitability but instead participate in diverse cellular processes including neurodevelopment, metabolism, and synaptic organization (31). Most current ASMs target voltage-gated ion channels, enhance GABAergic signaling, and modulate synaptic transmission to counteract neuronal excitability (34). Genetic heterogeneity in epilepsy has highlighted the need for systematic approaches to identifying novel pathways that may expand beyond conventional mechanisms.

In this study, we identified 2,496 epilepsy associated genes, analyzed their functional characteristics, and identified perturbed molecular clusters by diffusing gene-level evidence scores in an integrated brain-filtered network from WP. We sought to find the converging pathways underlying genetic heterogeneity in a human pathway-based network. The identified subnetworks are associated with neuronal development, membrane excitability, synaptic function, and cellular regulation. The subnetworks were highly enriched in ASM targets and interconnected by 44 first neighbors. The pathways represented by these subnetworks and their connecting nodes may offer opportunities for therapies. To illustrate this potential, we highlighted a subnetwork involved in cerebral cortex development and the connector CREB1, together with associated approved drugs identified in DrugBank.

Several research groups have demonstrated the use of network analysis to identify disease mechanisms and therapeutic targets. Lv et al. performed a topology-based network analysis using epilepsy risk genes and a PPI network to identify overlaps between epilepsy risk genes and ASM targets (35). Chu *et al*. also applied topology-based network analysis to genes associated with refractory epilepsies and identified 48 disease modules relevant to refractory epilepsies (36). These investigations examine the highest-degree nodes and the shortest-path distances to seed nodes. Although topology reveals valuable properties of the local network, it overlooks the global structure of the network (37).

Disease-related signals can be distributed across interacting genes, including those with weaker associations, making them difficult to identify through individual gene analysis. Diffusion-based algorithms model the perturbation signal flow by selecting nodes with initial weights and diffusing the weight through all connections (15). None of the 189 nodes identified by HotNet2 were ranked among the top 100 nodes by degree. This showed that the selected subnetworks were not dominated by highly connected genes. However, when expanding the subnetworks with their first-order neighbors, we found six out of ten top degree-ranked nodes connecting the subnetworks. This suggests that high-degree nodes may link disease-associated modules even though they did not eventually carry high heat.

One major consideration of our approach was the use of an integrated human pathway network as the background for propagation. Some seed-based PPI network constructions expand the input gene set to include up to 2^nd^ neighboring proteins. (38,39). After integrating the network, interactions between neighboring proteins may be lost. Protein-only networks might also exclude metabolite interactions, which are relevant to epileptogenesis, as we found that 20% of classified epilepsy-associated gene products were metabolite interconversion enzymes. To overcome these limitations, we sought to build an integrated human pathway-derived interaction network from WP. In WP, the interactions are human-curated and annotated with interaction types (inhibition, stimulation…), but they are presented in isolated pathways. We unified the identifiers of entities and kept only brain-expressed gene products. Other resources offer complementary approaches to representing molecular relationships (40). WP covers 90% of GO terms by gene products (41). WP was considered sufficient for biological coverage and its machine-readability.

Another consideration is curating the EAGs. The necessity to include multiple resources to complement data coverage was demonstrated by the fact that only 35 genes overlapped across the four resources. Other efforts have been made to catalogue epilepsy genes (8,31,42). For example, Genes4Epilepsy catalogues only monogenic genes from clinical diagnostic providers and panels. Our broader phenotype-based search aimed to capture additional associations. This approach increases coverage, although the resulting associations vary in evidence strength and should not be interpreted as established causal relationships. It is worth mentioning that from the HotNet2 output subnetwork components, we did not find genes that were not previously associated with epilepsy.

The ten subnetworks included in the visualization present converging subnetworks in genetic epilepsy. The development process is presented in S1 and S4. This finding is consistent with results of previous studies. Heyne *et al*. found genetic overlaps between de novo variants in neurodevelopmental diseases with and without epilepsy (43). S6 and S10 were associated with synaptic functions. The dysfunction of presynaptic and postsynaptic components has been well described, and they are targeted for ASM drug development (44), such as levetiracetam targeting SV2A (synaptic vesicle glycoprotein 2A) (45). TOR signaling (S5), a central regulator of cell growth and metabolism, is critical for epileptogenesis (46,47). Everolimus, an mTOR inhibitor, has been approved for add-on treatment for tuberous sclerosis complex-associated focal-onset seizures (48).

The resulting subnetworks not only show and identify the established converging mechanisms in genetic epilepsy but also provide potential targets. Drugs targeting one component of a subnetwork may also have therapeutic potential for patients with pathogenic variants in other genes within the subnetwork. As shown in **Fig 4**, we showed that from a pathway of interest, we can add prior knowledge of drug-target interactions. We showed in S4 (*cerebral cortex development*) that the connector (CREB1) connects most of the subnetworks. Among the three drugs targeting TUBA1A, a component from S4, albendazole plus dexamethasone reduced the number of seizure occurrences with generalization in 120 patients with living cysticerci in the brain and seizures (49). However, that trial addressed seizures caused by a parasitic brain infection, not genetic epilepsy, and the benefit could be attributed to killing the cysts. In another clinical trial involving patients with generalized tonic-clonic seizures, the 33 patients who received naloxone showed a shorter time to postictal mobility recovery than historical controls (50). Although this finding does not establish seizure-reducing efficacy or validate CREB1 as a therapeutic target, it supports the investigation of the effects and underlying mechanisms of naloxone.

## CONCLUSION

This study demonstrated that network-based analysis of epilepsy-associated genes can reveal functional molecular modules with translational relevance. By integrating disease-associated gene curation, interaction data, and heat diffusion clustering, we identified candidate targets and created a drug repurposing workflow for genetic epilepsy. The study provides a rational framework for generating hypotheses for future drug discovery in genetic epilepsy.

## DATA ACCESS

The code for data acquisition and processing is available in the GitHub repository at https://github.com/YuanyuanDuan1/Epilepsy-Interaction-Networks.

## AUTHOR CONTRIBUTIONS

Y.D. designed the work, performed the acquisition, analysis, and interpretation of data, and wrote the first draft of the manuscript. F.E. initially conceptualized the project and provided close supervision. F.E., A.V., R.P.W.R., R.J.V., and T.M.K. provided critical reviews of the manuscript and contributed to the final version.

## ACKNOWLEDGEMENTS

We thank Javier Millán Acosta and Martina Summer-Kutmon for supporting this work with advice on accessing WikiPathways data, and Denise Slenter and Tooba Abbassi-Daloii with advice on network construction.

## SUPPLEMENTARY MATERIALS

The supplementary materials are available at https://figshare.com/account/articles/34525656.

Supplementary File 1 Table S1. The entities retrieved from OMIM associated with epilepsy.

Supplementary File 1 Table S2. The entities retrieved from DisGeNET associated with epilepsy.

Supplementary File 1 Table S3. The entities retrieved from Orphadata associated with epilepsy.

Supplementary File 1 Table S4. The entities retrieved from ClinVar associated with epilepsy.

Supplementary File 1 Table S5. The mapped genes (EAGs) from four phenotype-genotype databases.

Supplementary File 1 Table S6. The significant GO terms and pathways of EAGs.

Supplementary File 1 Table S7. The complete edge lists of WikiPathways.

Supplementary File 1 Table S8. HotNet2 subnetwork outputs at each candidate δ threshold.

Supplementary File 1 Table S9. The 44 nodes that connect the subnetworks.

Supplementary File 1 Table S10. Antiseizure medications and their targets from Open Targets.

Supplementary File 2 Fig S1. The mapping to Ensembl IDs from the retrieved entities from OMIM, DisGeNET, Orphanet, and ClinVar.

Supplementary File 2 Fig S2. The top ten most significantly enriched GO MF(A), CC(B) and WP(C) ranked by adjusted p-values.

Supplementary File 2 Fig S3. All interaction counts from human pathways in WP.

Supplementary File 2 Fig S4. Comparison of weighting schemes using Jaccard similarity at the highest δ.

Supplementary File 2 Fig S5. Alluvial plot showing changes in subnetwork composition across four δ thresholds from HotNet2.

Supplementary File 2 Fig S6. The observed and expected subnetwork numbers in the WP-derived network at the highest δ.

